# Modelling Microbiome Association with Host Phenotypes Using a Bayesian Dirichlet Process Model

**DOI:** 10.1101/2024.10.08.617289

**Authors:** Denis Awany, Emile R. Chimusa

**Affiliations:** Institute of Infectious Disease and Molecular Medicine and Division of Immunology, Department of Pathology, University of Cape Town, Anzio Road, Cape Town, 7925, Western Cape, South Africa; Department of Applied Sciences, Northumbria University, Sutherland Building, Northumberland Road, Newcastle, United Kingdom

## Abstract

Dysbiosis in the human gut microbiome has been shown to be intimately involved in the pathogenesis of a wide range of communicable and non-communicable diseases. As microbiome wide association study becomes the workhorse for identifying association between microbial taxa and human diseases/traits, proper modelling of microbial taxa abundances is critical. In particular, statistical frameworks need to effectively model correlation among microbial taxa as well as latent heterogeneity across samples. Here, a Bayesian method using the Dirichlet process random effects model is devised for microbiome association study. The proposed method uses a weighted combination of phylogenetic and radial basis function kernels to model taxa effects, and a non-parametrically modelled latent variable to model latent heterogeneity among samples. Using simulated and real microbiome datasets, it is shown that the method has high statistical power for association inference.

**Software:** The R codes to implement the method has been incorporated into a script phyloDPM.R, and is available online at https://github.com/AwanyDenis/phyloDPM.

## 1. Introduction

The discovery of the association between the microbiome and multiple human phenotypes has spurred interest in microbiome research to identify biological and environmental factors that are associated with variation in microbiome composition, and to establish the link between microbiome and human clinical outcome (phenotypes).

To investigate the association between microbiome and host’s clinical outcome, microbiome samples are collected, genomic DNA are isolated, and then sequenced using next-generation sequencing (NGS) machines. Two general approaches are followed: shotgun metagenomic sequencing and gene-targeted sequencing. Shotgun metagenomic approach involves an unrestricted sequencing of the genome of all microorganisms present in a sample. The 16S rRNA method is, however, routinely used due to its relatively low cost. With this method, after sequencing and pre-processing, the 16S sequences are usually clustered into operational taxonomic units (OTUs) based on similarity, such as 97% similarity level, which are then considered to fairly represent species members, or amplicon sequence variants (ASVs).

The resulting species-level dataset has several intrinsic properties, including (1) over-dispersion with a large proportion of the taxa having zero or extremely low abundance across the samples; possibly due to sequencing error, the limit of the detection of sequencing machines or under-sampling for rare taxa, (2) complex correlation structure due to the phylogenetic tree structure imposed on the taxa. Thirdly, the individual-level samples exhibit marked variability in sequencing depth (library size), and (3) phylogenetic linkage, where it has been observed that phylogenetically-close taxa tend to have similar biological functions; the implication of which is that a given microbiome-associated host’s phenotype (clinical state) may result from the joint effect of multiple ‘clusters’ of taxa at a given taxonomic rank [1, 2]. Accordingly, leveraging the phylogenetic tree structure in analysis is critical for improved inference [3].

In this work, a Bayesian framework (herein named ‘phyloDPM’) for identifying OTU-specific association with a given host’s phenotype is proposed. The proposed model uses a weighted combination of the phylogenetic- and radial basis function-defined similarity kernels to model the covariance structure of taxa effects, and a Dirichlet process (DP) prior to model latent heterogeneity across samples. As the phylogenetic tree itself can be noisy or uninformative with respect to the phenotype under study [4], using composite kernels leads to improved inference. Meanwhile, by modelling the latent variable non-parametrically, the restrictive distributional assumption often made on such random effects is avoided and, instead, the distribution for the parameters is allowed to adapt to the data. In this way, the model remains robust under a wide range of sample features, including multi-modality, heavy tailedness, and skewness; all of which would otherwise result in false inference about the true magnitude of the estimated effects if the true data distribution differs from the assumed model [5]. The performance of the method is illustrated through application to simulated dataset, and to two real microbiome datasets of schizophrenia, and atherosclerosis diseases.

## 2. Materials and Method

### 2.1. Model Specification

Consider a microbiome study in which a total of *p* OTUs have been profiled from *n* individuals. Suppose also that the phenotype of interest has been recorded for each individual.The OTU data can be represented in an *n* × *p* matrix, with rows corresponding to the samples and columns corresponding to the OTUs. A characteristic feature of such data is that the total number of OTUs (which is usually in hundreds or thousands) is much larger than the sample size *n* (which is usually in tens or hundreds). In addition, there is intrinsic correlation among the OTUs, imposed by the phylogenetic tree structure. The phylogenetic tree structure can be specified by a *p* × *p* matrix *D* of pairwise patristic distances between OTUs; this distance matrix being independent of the observations in the *n* × *p* data matrix.

We are interested in modelling the association between the OTUs and the phenotype (outcome) of the host. To cast this into a mathematical formulation, let **y** = (*y*_1_, *y*_2_, · · ·, *y*_*n*_), be the binary response of the *n* individuals; *X*, a *n* × *q* matrix of covariates (e.g gender, age, etc); and *Z*, a *n* × *p* matrix of normalized taxa abundance associated with the observations. Let *ψ* = (*ψ*_1_, *ψ*_2_, · · ·, *ψ*_*n*_), be a latent variable that accounts for subject-specific deviation from the underlying model. The association of the OTUs with the phenotype, for individual *i*, is then modelled by the logistic random effects model given by

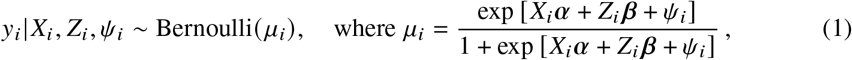

*α* = (*α*_1_, · · ·, *α*_*q*_), is a *q*-dimensional vector of covariate effects including an intercept, and *β* = (*β*_1_, · · ·, *β*_*p*_), is a *p*-dimensional vector of OTU effects. We seek two primary aims: (1) to account for multicollinearity among bacterial taxa at various phylogenetic depths, by using a weighted combination of co-informative kernels, the phylogenetic and radial basis function (RBF) kernels, to define the covariance structure for taxa effects, and (2) to account for latent structuring (similarity) across samples, by modelling the latent variable *ψ* as realizations from a Dirichlet process (DP).

#### 2.1.1. Phylogeny-based Correlation among OTUs

To incorporate the phylogenetic tree information into the analysis, we base on the patristic distance matrix *D* to define the correlation structure among the OTUs. Using the trait evolution model [6], the correlation of traits between OTU *j* and OTU *k*, which we specify as *C*_*jk*_, can be modelled using the following exponential function

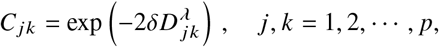

where *D* _*jk*_ is the patristic distance between OTU *j* and OTU *k*, and *δ* ∈ [0, ∞) is an evolutionary rate parameter.

Considering *δ*, a *δ* = 0 means that there is no evolution. In this case, *C*_*jk*_ = 1 ∀ *j, k* which implies that all traits are identical with maximum phylogenetic relatedness. Meanwhile, *δ* → ∞ means extremely rapid evolution. The result of this is that *C*_*jk*_ → 0 ∀ *j* ≠ *k*, with the implication that the traits have no phylogenetic relationship. From this observation, *δ* can also be viewed as an OTU dependency measure, by which *δ* = 0 means all the OTUs are dependent (intricately linked) and *δ* → ∞ all the OTUs are independent. As a corollary, the magnitude of *δ* also determines the OTU phylogenetic relatedness at the various taxonomic ranks. Denote this phylogeny-based correlation matrix by *C*_*phy*_.

#### 2.1.2. Radial basis function-based Correlation among OTUs

The relationship matrix, *C*_*phy*_, defined above captures signals from OTU effects, permitting a parametric analysis and the interpretation with respect to the underlying genealogical architecture of microbial taxa. It is also desirable to invoke a similarity measure that captures characteristics of the data, regardless of the underlying phylogenetic relationship. A radial basis function (RBF) kernel, as similarity matrix, allows interpretation of classical relatedness as spatial distances between taxa.

Treating **z** as continuous, then, without loss of generality, the spatial distance between OTUs *j* and *k* is given by

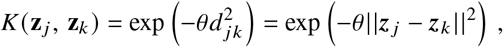

where *θ* ∈ ℝ^+^ is a scaling parameter that determines the overall scale on which the abundance varies across taxa. For very rapidly varying abundance, it is small, and for slowly varying abundance it is large. The smaller the Euclidean distance between two OTUs, the stronger the similarity exists between them. The kernel matrix serves as the covariance matrix for modelling OTU effects; denote this kernel matrix by *C*_*rb f*_.

Since both *C*_*phy*_ and *C*_*rb f*_ are Gram matrices [7], they are positive definite. The composite covariance matrix, *C*_*T*_, is set up as

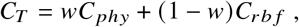

where *w* ∈ [0, 1] is a weighting parameter. In this exposition, the correlation is modelled as a linear combination of a phylogenetically- and RBF-based similarities. *w*, which is constrained to ensure positive semi-definiteness of the composite kernel, controls the relative contribution of the phylogenetic and radial basis function kernels to defining the correlation structure among microbial taxa. When *w* = 1, the implicit assumption is that the phylogenetic tree completely specifies correlation among the microbial taxa and the RBF kernel is uninformative. A *w* = 0, however, implies the RBF sufficiently describes the microbial correlation structure and the phylogenetic tree is entirely uninformative. When *w* is selected as 0.5, the radial basis function and the phylogenetic structure are allowed to contribute equally to define the correlation structure.

The joint covariance, *C*, is used to regularize taxa effects. Specifically, let the coefficient vector *β* follow a multivariate normal distribution, that is,

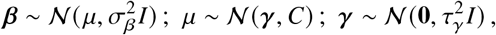

where *I* is the *p* × *p* identity matrix, 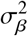 and 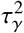 are unknown variance components, and the vector *μ* is the mean of the Gaussian distribution for the main effects of taxa.

### 2.2. Dirichlet Process for the Random Effects ψ

We model the heterogeneity in OTU abundance across individuals as random effects that follow a Dirichlet Process (DP). The DP, specified as a prior in a Bayesian setting, has several appealing features, including incorporating multimodality, expressing our uncertainty about the form of the underlying random effects, and robustness to errors in model specification.

Let each random effect *ψ*_*i*_ be a realization from a general distribution that follows a DP, and suppose further that the base distribution, *G*_0_(*B*, is Gaussian with concentration (precision) parameter *m*(> 0). Then,

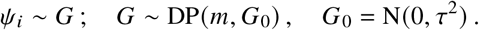

The predictive distribution of *ψ* = (*ψ*_1_, *ψ*_2_, · · ·, *ψ*_*n*_) is

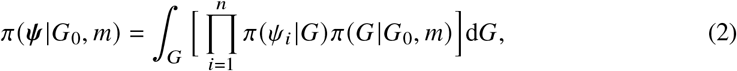

which can be decomposed as

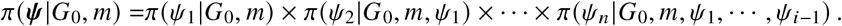

Following [8], we have *ψ*_1_|*G*_0_, *m* ∼ *G*_0_, and the successive conditional distributions are

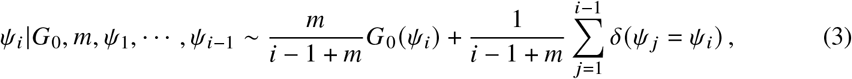

where *δ*(·) is the usual Kronecker delta function. From this formulation, it can be seen that there is a non-zero probability that there will be ties among the realizations. The implication of this is that the DP generates clusters among the observations. The clustering assigns different base distribution parameters across groups and the same parameter within groups. Thus, *ψ*_*i*_ values are independent and identically distributed (i.i.d) only if they are within the same group. This, which is in contrast to the usual parametric modelling strategy that presumes that the random effects are all i.i.d, represents a more general method.

### 2.3. Model Likelihood

The joint distribution of the observed data in Eqn (1) can be expressed in a general form as

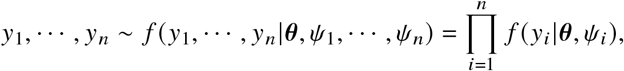

where θ is a vector of all except the DP model parameters. The likelihood function, *L* (θ |**y**), can then be obtained, by definition, by integrating over the random effects. Therefore

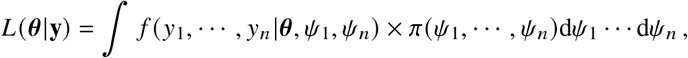

where *π*(*ψ*_1_, · · ·, *ψ*_*n*_) = *π*(*ψ*) is as defined in Eqn (2).

The clustering in the DP model leads to a succinct mathematical formalism in which the observations can be represented by a binary matrix. Following [5, 9], let *A* be a *n* × *k* binary matrix associated with a particular partition of the sample of size *n* into *k* groups, *k* = 1, · · ·, *n*. A partition as such is a cluster or, in a more appropriate terminology, a ‘subcluster’ *C*, since the grouping is done non-parametrically and the resulting clusters may differ from that determined by some substantive criteria. Each row of *A* has all zeros except for a 1 in one position indicating the group the observation belongs, whereas the column sums of *A* represent the number of observations in each of *k* groups. Therefore, the *n*-dimensional vector *ψ* can be represented by a *k* (≤ *n*)-dimensional vector ***η***. Then *ψ*_*i*_ = *η*_*j*_ so that ***ψ* = *Aη***, where ***η*** ∼ *N*_*k*_ (0, *τ*^2^*I*_*k*_).

The likelihood function becomes

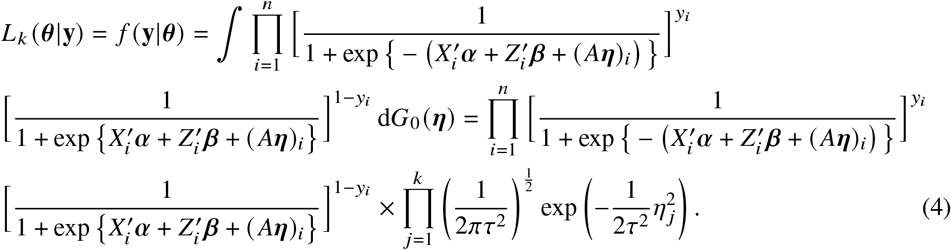

To complete the Bayesian model, we specify the following priors on the other model parameters:

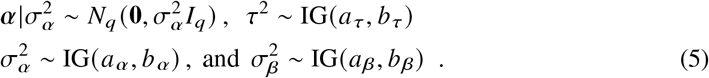

### 2.4. Bayesian Inference of the Parameters

Combining the likelihood in Eqn 4 with the priors in Eq 5 gives the following joint posterior distribution (details in Supplementary material).

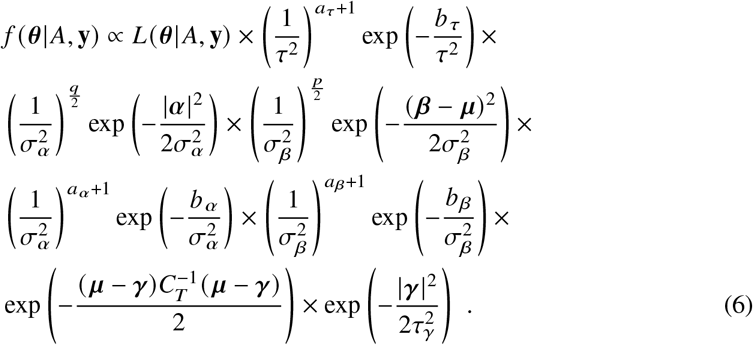

The complexity of the dependency structure of the model, imposed by the DP components, makes direct Markov Chain Monte Carlo (MCMC) sampling of the lower-level parameters using the Gibbs sampler impossible to implement, because the resulting conditional distributions assume awkward forms. In this case, one may be tempted to turn to the Metropolis-Hastings algorithms. However, these algorithms can be difficult to set up for such complex functional forms, and may require ‘tuning’ to achieve satisfactory performance. Moreover, even though the ‘black box’ random variate generation techniques such as the ratio-of-uniforms method may seem an alternative, their use can be daunting since they also require tuning to achieve reliable and efficient sampling.

To circumvent these difficulties, we invoke the auxiliary variable (*slice* sampling) method [10]. The general strategy here is to augment the parameter space with a set of convenient random (auxiliary) variables that keeps the marginal posterior distribution of interest unchanged but converts all of the full conditionals into a set of easily sampled ‘standard’ full conditional distributions. Additionally, the technique also improves sampling efficiency [11, 12]. By this, the slice-Gibbs sampling can be used to sample the lower-level parameters. Meanwhile, the higher-level parameters can be directly sampled using the Gibbs sampler applied to their respective full-conditional distributions.

### 2.5. Full Conditional Distributions

Estimates of model parameters are obtained by sampling from the full conditional distributions. For notational convenience, let ***θ*** be the vector of all model parameters and ***θ***_−*p*_ be the vector of all these parameters except *p*.

Then (Supplementary material), we obtain the following full conditional distributions for the parameters:

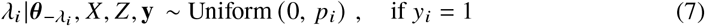

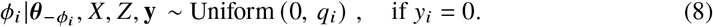

The full conditional distribution for *α* becomes the truncated Gaussian distribution:

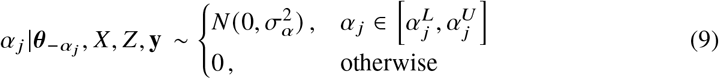

where *α*_*j*_ ; *j* ∈ {1, · · ·, *q*}, and *α*_*j*_ ; *j* ∈ {1, · · ·, *q*} have the following full conditional distributions

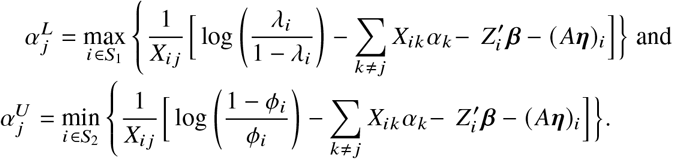

The full conditional distribution for *β*_*j*_ ; *j* ∈ {1, · · ·, *p*} is

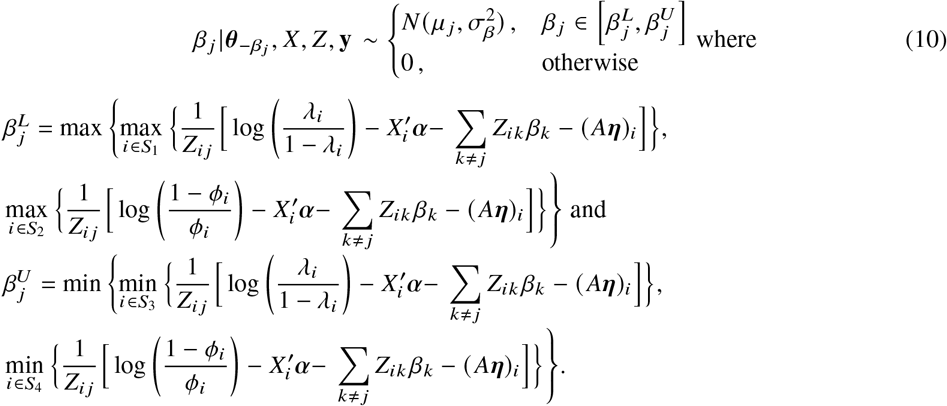

The full conditional distribution for *η*_*k*_ ; *k* ∈ {1, · · ·, *K*} is

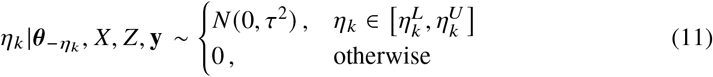

where 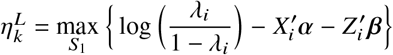, and 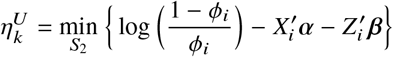.

The full conditional distribution for *μ*_*j*_ is:

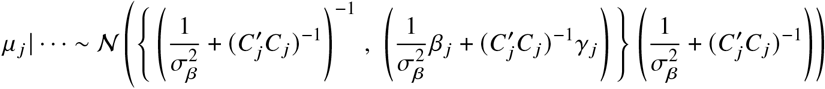

The full conditional distribution for ***γ*** is

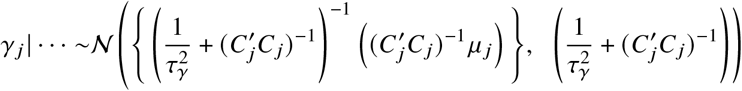

Finally, the full conditional distribution for *τ*^2^, 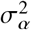, and 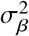 are, respectively:

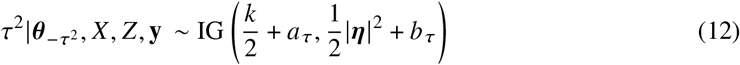

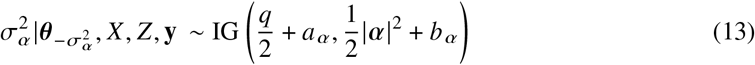

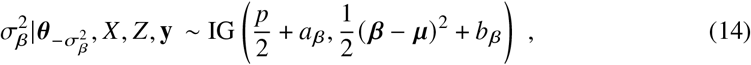

where IG(·, ·) denotes the inverse Gamma distribution.

### 2.6. Generation of Sub-clusters Matrix A

The sub-clusters matrix *A* is generated by iteratively sampling from a Multinomial distribution as:

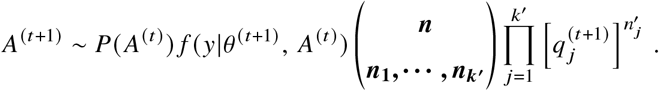

In the sampling, columns with zero column sums are deleted resulting in a *n* × *k* matrix; the deletion of the zero sum columns is a marginalization of the multinomial distribution.

### 2.7 Estimation of Precision Parameter m

The precision parameter *m*, unknown before hand, plays pivotal role in the distribution of ***ψ*** as it determines the number of unique clusters. Adopting [13]’s sampling scheme, we estimate *m* by first sampling an auxiliary variable *v* is sampled from a Beta distribution, i.e. *v*|*m, k*, · · · ∼ Beta(*m* + 1, *n*), and then sampling *m* from the mixture:

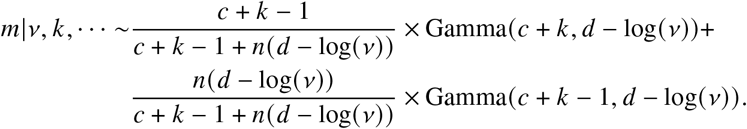

In this way, *m* can be iteratively updated within the general Gibbs sampling framework, conditional on the other model parameters.

## 3. APPLICATION

### 3.1. Simulation Studies

For this, counts of OTU were simulated by sampling from a Dirichlet Multinomial (DM) distribution. In order to make the simulation more realistic, the parameters of the DM distribution (mean and dispersion) were estimated from real microbiome data of human upper respiratory microbiome in the study by [14]. This dataset contains the counts of 856 OTUs from 60 samples. Also available from this dataset is the phylogenetic tree structure for these OTUs.

The *p* ∈ {200, 400, 800} most abundant OTUs from this dataset were used for parameter estimation, and, accordingly, *p* OTUs were subsequently simulated for *n* = 500 individuals. The simulated OTU counts data closely mimic real data (**Supp Fig. 1**).

The resulting OTU count data for each sample was normalized by the total read count for that sample, where the total read count for a sample was obtained by sampling from a negative binomial distribution with mean 5000 and dispersion 25, which reflects typical sequencing depths in empirical microbiome studies [4].

To simulate a phylogeny-informative scenario, the partitioning-around-medoids (PAM) algorithm was used to partition the *p* OTUs into clusters into 50 different sizes, based on the patristic distances between them. Then, one cluster with the highest abundance was chosen and ten most-abundant OTUs from this cluster were selected to be associated to the phenotype.

We introduce cluster effects in the simulation by simulating random effects, *ψ*, that follow a Dirichlet process. For each individual, *ψ*_*i*_ ∼ *G*, and *G* ∼ DP(*m, G*_0_). We let *G*_0_ ∼ *N* (0, 1), and to set a value for *m*, we arbitrarily fix the number of subclusters, *K*, to 40 (8% of the sample size), and make use of the mathematical relationship between *K* and *m*, which is 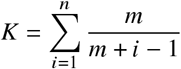, to obtain *m*. This yields *m* = 24.21. With *G*_0_ and *m*, we use the Polya-urn model in (3) to simulate *ψ*_*i*_, (*i* = 1, · · ·, *n*).

Finally, the phenotype, *y*_*i*_, (*i* = 1, · · ·, *n*) is simulated from the model *y*_*i*_ = Bin(*p*_*i*_); 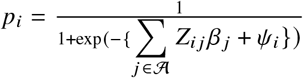, where 𝒜 is the set containing the associated OTUs, and the effect size *β*_*j*_ is varied between 0.1 and 2.0, which corresponds to typical effect sizes observed in microbiome studies [15, 16].

#### 3.1.1. Simulation Results and Discussion

We first conducted test runs to check the convergence property of the chain, and found that in each case the chain mixes properly within 5000 iterations. Convergence of the chains were assessed by visual inspection using trace plot, auto-correlation function plot and cumulative quantile plot, as well as formal diagnostic test using Geweke disgnostic [17]. All these showed no evidence of non-convergence. We observe from these plots that the chain moves quite quickly over the possible values of the parameters, moving directly to the center of the target posterior distribution. Following this, the Markov chain was run for 50000 iterations from where the first 5000 was discarded as burn-in, and, to remove the autocorrelation, a lag of 10 was considered. The resulting effective samples were the ones finally used for posterior inference (**Table 1**). Because we know the true values of the simulated effects, we can now assess the model’s performance. Optimal hyper-parameters for the model were obtained by sensitivity analysis (**Supp Fig. 2**).

**Table 1.**
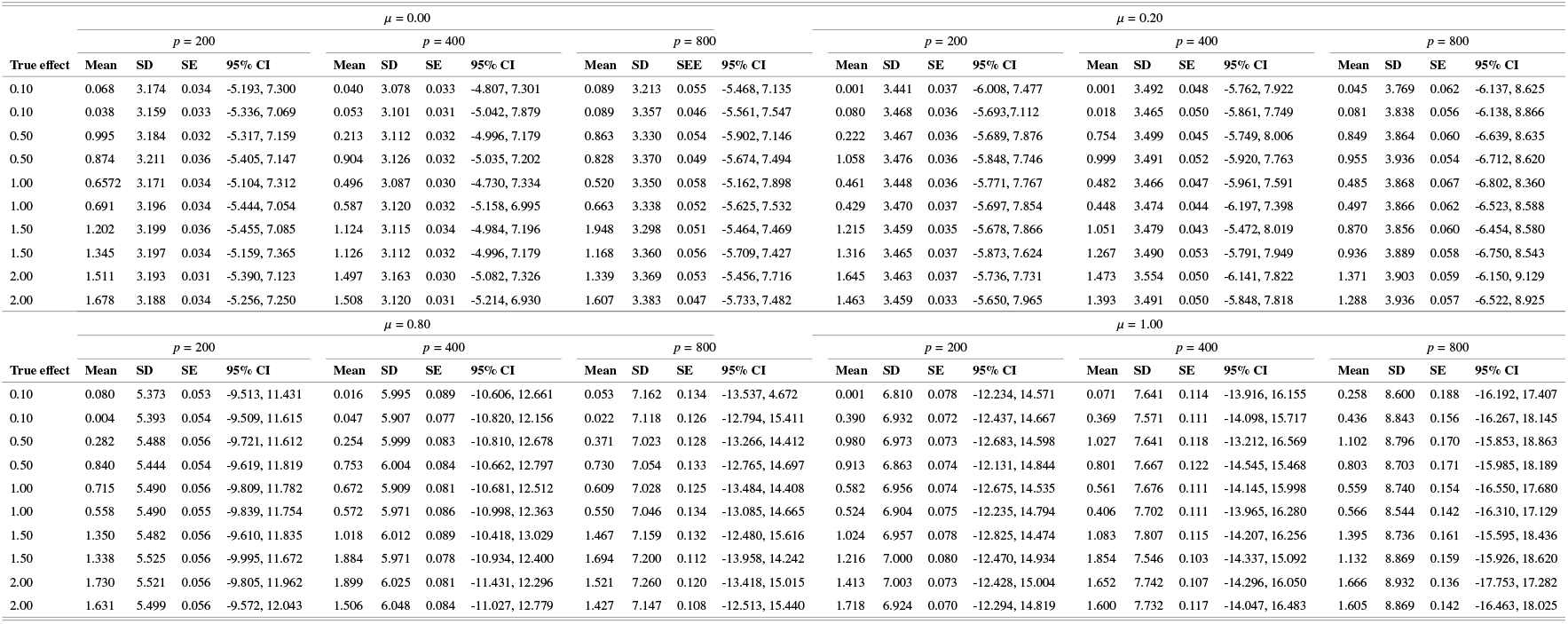
Posterior summary statistics produced by fitting the model to simulated data. Posterior mean (Mean), standard deviation (SD), standard error (SE), and 95% credible interval (CI) of the estimated effects.

We find that the proposed method performs satisfactorily well under all the scenarios considered (**Table 1**). Most noticeably, the method performs well when comparing estimates of *β*_*j*_ with the true values. It can be seen that the 95% credible intervals cover the true parameter value for all the model parameters (OTU effects), standard errors are very low for virtually all coefficients.

We now turn to the impact of the weighting parameter *μ* (that determines the relative importance of the Kernel- and the phylogeny-based similarity) and the number of OTUs *p*. It can be seen that both *μ* and *p* had discernible influence on empirical performance, for a fixed sample size. For *μ* = 0, the standard errors are roughly similar for *p* = 200 and *p* = 400. However, as *p* increases from 200 to 800, the standard errors increase as *p* becomes large (*p* = 800). Similar tends are observed for *μ* = 0.2, 0.8 and 1.0; the accuracy diminishes as *p* increases.

Meanwhile, for any fixed value of *p*, the standard errors are very low for small values of *μ* (**Supp Fig. 3**). One possible explanation for this is that the phylogenetic tree is noisy or uninformative - an issue that has been noted for phylogenetic tree usage in microbiome analysis [3] - and therefore as *μ* is increased, by which the phylogeny-defined similarity becomes more favored than the radial basis function one, the noise introduced in the data reduces model fit.

### 3.2. Real Data Studies

#### 3.2.1. Application to Schizophrenia

For empirical evaluation of performance in identifying associated bacterial taxa, we applied the method to a real microbiome data, comparing the gut microbiome composition between persons with chronic schizophrenia and non-psychiatric individuals. The microbiome abundance data from 16S rRNA sequencing were obtained from 25 chronic schizophrenic and 25 demographically matched non-psychiatric subjects. The dataset is available for download from Qiita (https://qiita.ucsd.edu; study ID 11710). Sequence reads were processed using QIIME2 [18]; raw sequencing results demultiplexed and unique microbial OTUs identified using Deblur algorithm [19], and the resulting output feature table rarefied to 7,905 sequences per sample. A total of 3,564 OTUs were obtained (Supplementary material). The resulting OTU (‘species’ level) data were agglomerated to, and analysed at, the class taxonomic level. The corresponding phylogenetic tree was accordingly prunned to contain only these taxa.

The MCMC algorithm was then run for 50,000 iterations. A burn-in period of 10,000 was considered and the remaining effective samples thinned by a factor of 10 before performing posterior inference. As a validation, we also analyse the same data using the standard generalized linear model (GLM), which essentially is a Bayesian probit model with highly diffuse priors. As the standard generalized linear model accounts for neither sample clustering nor correlation among the predictors (with no phylogenetic tree incorporated), our aim was not to view it as an alternative competing model, as this would certainly be biased, but rather to demonstrate the trade-offs and performance of schemes that incorporate phylogenetic information and/or account for samples clusterings with those that do not.

The results are summarized in **Table 2**, showing the posterior means (i.e., the model coefficients) for the phyloDPM and the GLM. From it, we see that the samples exhibit substantial clustering, as indicated by the precision parameter, *m*, of the DP and the number of clusters, *K* (= 22), which is less than half the sample size *n*(= 49). We observe that, apart from the sign of the estimated effects being consistent across the two models, the phyloDPM provides a more conservative estimate of coefficients. That is, by this, the statistically reliable estimate of effects are the values actually obtained (low or high) and not those that ought to have been obtained (high or low).

**Table 2.** Summary of parameter estimates for the schizophrenia data: posterior mean, standard error (SE) and 95% credible interval (CI) for microbial taxa effects. Results averaged for 30 replications.

| Item | phyloDPM |  |  |  | GLM |  |  |  |
| --- | --- | --- | --- | --- | --- | --- | --- | --- |
|  | Mean | SE | 95% CI |  | Mean | SE | 95% CI |  |
| Intercept | 0.062 | 0.016 | -0.777 | 0.890 | 3.773 | 4.844 | -5.721 | 13.268 |
| *Clostridia* | -0.361 | 0.013 | -0.797 | 0.040 | -0.326 | 0.230 | -0.777 | 0.126 |
| *Bacilli* | -0.052 | 0.008 | -0.360 | -0.262 | -0.086 | 0.133 | -0.347 | 0.174 |
| *Bacteroidia* | 0.071 | 0.010 | -0.063 | 0.239 | 0.026 | 0.046 | -0.065 | 0.117 |
| *Actinobacteria* | 0.063 | 0.004 | -0.107 | 0.234 | 0.042 | 0.078 | -0.112 | 0.196 |
| *Coriobacteriia* | 0.433 | 0.022 | 0.074 | 1.014 | 0.313 | 0.166 | -0.013 | 0.640 |
| *Deltaproteobacteria* | 0.028 | 0.010 | -0.358 | 0.327 | 0.019 | 0.178 | -0.330 | 0.369 |
| *Erysipelotrichi* | -0.030 | 0.007 | -0.222 | 0.176 | -0.034 | 0.074 | -0.179 | 0.110 |
| *Gammaproteobacteria* | 0.033 | 0.001 | -0.021 | 0.092 | 0.029 | 0.024 | -0.019 | 0.076 |
| *m* | 17.750 | 0.027 | 9.323 | 27.454 | - | - | - | - |
| *K* | 22.124 | 0.176 | 0.000 | 29.000 | - | - | - | - |

To put these results into context, it is observed that microbial taxa of the class *Coriobacteriia* tends to have a higher probability of association with the disease condition. Microbial taxa of the class *Coriobacteriia* are Gram-positive bacteria within the *Actinobacteria* phylum. In previous studies [20, 21], *Actinobacteria* has been shown to be relatively enriched in the gut microbiome of smokers compared with that of non-smokers, and that higher abundance of the *Actinobacteria* phylum was significantly associated with schizophrenia. The result herein obtained is therefore consistent with literature. Meanwhile, the result also suggest that disease risk reduces in presence of taxa such as *Clostridia, Bacilli, Bacteroidia*, and *Erysipelotrichi*; consistent with previous observations [21]. However, in contrast with earlier findings, the abundance of *Deltaproteobacteria* have not be reported to be repressed in schizophrenic subjects, as the result here indicate.

#### 3.2.2. Application to Atherosclerosis

As a final illustration of the performance of the phyloDPM model, we further analyze atheroscle-rosis data with 4623 OTUs from 73 samples [Qiita, https://qiita.ucsd.edu; study ID 349]. Following the pre-processing procedure, a total of 652 OTUs remained across 72 samples. The underlying phylogenetic tree was also subsequently prunned to these remaining OTUs.

The top 50 taxa showing strongest association are shown in **Table 3**. In general, the posterior means (i.e, the coefficients) for the effects of these OTUs range ∼ 0.04 − 0.12. These effect sizes are consistent with typical values reported in microbiome studies [15, 16]. The standard errors are small for virtually all the coefficients, showing that the method maintains its satisfactory performance. However, the credible intervals of the parameters are fairly large; possibly attributable to the outliers that unduly influence model parameter estimates and distort credible intervals, tending to make them wide.

**Table 3.** Summary of parameter estimates for the Atherosclerosis data: posterior mean, standard deviation (SD), standard error (SE) and 95% credible interval (CI) for OTU effects. Also shown are the corresponding species and classes to which the various OTUs belong. The (-) on the species column indicates the species information is unavailable.

| OTU | Species | Class | Mean | SD | SE | 95% CI | OTU | Species | Class | Mean | SD | SE | 95% CI |
| --- | --- | --- | --- | --- | --- | --- | --- | --- | --- | --- | --- | --- | --- |
| OTU274 | - | Clostridia | 0.071 | 4.203 | 0.056 | -8.261 8.262 | OTU334 | - | Clostridia | 0.048 | 4.173 | 0.057 | -8.095 8.318 |
| OTU607 | - | Bacilli | 0.120 | 4.220 | 0.059 | -8.084 8.513 | OTU616 | - | Bacilli | 0.057 | 4.163 | 0.055 | -7.947 8.402 |
| OTU424 | Nanceiensis | Bacteroidia | 0.064 | 4.191 | 0.057 | -8.165 8.357 | OTU216 | - | Bacteroidia | 0.043 | 4.193 | 0.055 | -8.212 8.296 |
| OTU76 | - | Clostridia | 0.094 | 4.235 | 0.062 | -8.096 8.531 | OTU249 | - | Gammaproteobacteria | 0.052 | 4.179 | 0.056 | -8.230 8.242 |
| OTU347 | Sobrinus | Bacilli | 0.074 | 4.173 | 0.054 | -8.128 8.295 | OTU204 | hiranonis | Clostridia | 0.050 | 4.178 | 0.058 | -8.280 8.158 |
| OTU6 | - | Clostridia | 0.061 | 4.189 | 0.054 | -8.178 8.281 | OTU453 | - | Bacilli | 0.042 | 4.160 | 0.056 | -8.167 8.185 |
| OTU458 | - | Clostridia | 0.060 | 4.154 | 0.056 | -8.222 8.118 | OTU372 | - | Bacteroidia | 0.041 | 4.170 | 0.056 | -8.134 8.284 |
| OTU179 | - | Clostridia | 0.059 | 4.198 | 0.057 | -8.273 8.318 | OTU32 | - | Clostridia | 0.041 | 4.189 | 0.054 | -8.248 8.255 |
| OTU68 | - | Clostridia | 0.066 | 4.178 | 0.054 | -8.313 8.132 | OTU426 | - | Clostridia | 0.046 | 4.189 | 0.055 | -8.151 8.302 |
| OTU15 | Gnavus | Clostridia | 0.063 | 4.194 | 0.057 | -8.239 8.265 | OTU184 | - | Bacilli | 0.043 | 4.191 | 0.056 | -8.148 8.425 |
| OTU218 | - | Clostridia | 0.065 | 4.199 | 0.051 | -8.057 8.508 | OTU155 | - | Clostridia | 0.043 | 4.171 | 0.057 | -8.291 8.177 |
| OTU84 | - | Clostridia | 0.059 | 4.188 | 0.053 | -8.172 8.300 | OTU568 | - | Clostridia | 0.041 | 4.189 | 0.053 | -8.033 8.520 |
| OTU8 | Gnavus | Clostridia | 0.064 | 4.175 | 0.057 | -8.188 8.297 | OTU268 | Fragilis | Bacteroidia | 0.043 | 4.149 | 0.057 | -8.201 8.057 |
| OTU309 | - | Clostridia | 0.066 | 4.194 | 0.059 | -8.259 8.242 | OTU480 | - | Clostridia | 0.044 | 4.181 | 0.057 | -8.227 8.254 |
| OTU262 | - | Clostridia | 0.097 | 4.170 | 0.056 | -7.995 8.470 | OTU535 | - | Clostridia | 0.053 | 4.207 | 0.054 | -8.294 8.247 |
| OTU420 | - | Clostridia | 0.061 | 4.208 | 0.057 | -8.268 8.335 | OTU512 | Gnavus | Clostridia | 0.042 | 4.167 | 0.052 | -8.095 8.277 |
| OTU35 | - | Clostridia | 0.112 | 4.187 | 0.054 | -8.329 8.253 | OTU264 | - | Actinobacteria | 0.054 | 4.151 | 0.055 | -8.030 8.312 |
| OTU29 | - | Clostridia | 0.096 | 4.207 | 0.053 | -8.380 8.146 | OTU181 | - | Clostridia | 0.052 | 4.200 | 0.057 | -8.150 8.421 |
| OTU151 | - | Clostridia | 0.073 | 4.168 | 0.057 | -8.038 8.425 | OTU379 | - | TM7-3 | 0.050 | 4.181 | 0.055 | -8.213 8.270 |
| OTU105 | - | Actinobacteria | 0.060 | 4.180 | 0.054 | -8.001 8.459 | OTU392 | - | Betaproteobacteria | 0.041 | 4.207 | 0.056 | -8.184 8.357 |
| OTU552 | - | Clostridia | 0.062 | 4.232 | 0.056 | -8.137 8.617 | OTU124 | - | Clostridia | 0.042 | 4.177 | 0.053 | -8.178 8.339 |
| OTU465 | - | Clostridia | 0.061 | 4.189 | 0.054 | -8.208 8.260 | OTU73 | - | Clostridia | 0.042 | 4.194 | 0.059 | -8.289 8.210 |
| OTU136 | - | Bacilli | 0.059 | 4.172 | 0.057 | -8.156 8.317 | OTU19 | - | Bacteroidia | 0.041 | 4.195 | 0.056 | -8.011 8.563 |
| OTU128 | - | Betaproteobacteria | 0.067 | 4.184 | 0.053 | -8.156 8.275 | OTU324 | - | Bacilli | 0.046 | 4.207 | 0.057 | -8.248 8.461 |
| OTU41 | - | Actinobacteria | 0.058 | 4.173 | 0.057 | -8.048 8.400 | OTU339 | - | Gammaproteobacteria | 0.040 | 4.162 | 0.053 | -8.159 8.264 |
| OTU58 | - | Clostridia | 0.113 | 4.208 | 0.057 | -8.216 8.334 | OTU377 | - | Actinobacteria | 0.055 | 4.187 | 0.055 | -8.192 8.321 |
| OTU415 | - | Bacilli | 0.064 | 4.214 | 0.058 | -8.405 8.239 | OTU145 | - | Coriobacteriia | 0.043 | 4.173 | 0.058 | -8.222 8.237 |
| OTU137 | - | Clostridia | 0.066 | 4.154 | 0.054 | -8.254 8.162 | OTU635 | - | Actinobacteria | 0.045 | 4.176 | 0.056 | -8.350 8.142 |
| OTU531 | - | Clostridia | 0.105 | 4.163 | 0.054 | -8.128 8.288 | OTU331 | - | Gammaproteobacteria | 0.053 | 4.076 | 0.056 | -7.890 8.216 |
| OTU186 | - | Coriobacteriia | 0.065 | 4.197 | 0.058 | -8.274 8.290 | OTU437 | - | Bacilli | 0.044 | 4.162 | 0.055 | -8.180 8.246 |
| OTU450 | - | Clostridia | 0.091 | 4.154 | 0.057 | -8.089 8.241 | OTU196 | - | Clostridia | 0.051 | 4.149 | 0.058 | -8.216 8.139 |
| OTU387 | - | Clostridia | 0.060 | 4.191 | 0.058 | -8.156 8.383 | OTU633 | - | Clostridia | 0.041 | 4.205 | 0.056 | -8.364 8.239 |
| OTU251 | Dispar | Clostridia | 0.066 | 4.173 | 0.055 | -8.142 8.325 | OTU248 | - | Clostridia | 0.045 | 4.165 | 0.055 | -8.120 8.362 |
| OTU444 | Endodontalis | Bacteroidia | 0.073 | 4.179 | 0.059 | -8.230 8.267 | OTU266 | - | Fusobacteriia | 0.048 | 4.186 | 0.056 | -8.108 8.457 |
| OTU398 | - | Bacteroidia | 0.060 | 4.178 | 0.054 | -8.211 8.235 | OTU88 | - | Clostridia | 0.041 | 4.191 | 0.057 | -8.248 8.257 |
| OTU490 | - | Bacilli | 0.058 | 4.205 | 0.056 | -8.213 8.285 | OTU576 | - | Clostridia | 0.041 | 4.163 | 0.053 | -8.066 8.332 |
| OTU571 | - | Clostridia | 0.112 | 4.178 | 0.055 | -8.065 8.412 | OTU551 | - | Bacteroidia | 0.055 | 4.184 | 0.051 | -8.121 8.367 |
| OTU16 | - | Clostridia | 0.081 | 4.192 | 0.055 | -8.236 8.306 | OTU539 | - | Clostridia | 0.051 | 4.161 | 0.057 | -8.120 8.295 |
| OTU98 | - | Clostridia | 0.117 | 4.189 | 0.056 | -8.280 8.300 | OTU146 | Anginosus | Bacilli | 0.056 | 4.188 | 0.057 | -8.230 8.226 |
| OTU86 | - | Clostridia | 0.062 | 4.210 | 0.058 | -8.233 8.342 | OTU199 | - | Clostridia | 0.043 | 4.179 | 0.053 | -8.204 8.251 |
| OTU433 | - | Clostridia | 0.065 | 4.144 | 0.055 | -8.031 8.265 | OTU487 | - | Gammaproteobacteria | 0.044 | 4.184 | 0.057 | -8.185 8.207 |
| OTU99 | - | Fusobacteriia | 0.076 | 4.189 | 0.058 | -8.155 8.343 | OTU192 | - | Bacteroidia | 0.051 | 4.213 | 0.059 | -8.235 8.380 |
| OTU587 | - | Actinobacteria | 0.065 | 4.196 | 0.057 | -8.120 8.473 | OTU161 | - | Bacilli | 0.043 | 4.207 | 0.059 | -8.254 8.269 |
| OTU343 | - | Actinobacteria | 0.095 | 4.171 | 0.057 | -8.127 8.277 | OTU500 | - | Bacilli | 0.052 | 4.176 | 0.056 | -8.108 8.390 |
| OTU92 | - | Clostridia | 0.077 | 4.178 | 0.056 | -8.022 8.405 | OTU26 | - | Clostridia | 0.051 | 4.184 | 0.053 | -8.271 8.182 |
| OTU18 | - | Actinobacteria | 0.064 | 4.175 | 0.054 | -7.981 8.434 | OTU602 | - | Clostridia | 0.043 | 4.197 | 0.058 | -8.113 8.481 |
| OTU564 | - | Clostridia | 0.074 | 4.207 | 0.057 | -7.935 8.636 | OTU282 | - | Clostridia | 0.050 | 4.186 | 0.055 | -8.140 8.381 |
| OTU533 | - | Bacilli | 0.059 | 4.176 | 0.054 | -7.955 8.470 | OTU225 | - | Clostridia | 0.041 | 4.178 | 0.055 | -8.123 8.339 |
| OTU466 | - | Bacilli | 0.067 | 4.186 | 0.057 | -8.175 8.322 | OTU12 | - | Clostridia | 0.046 | 4.217 | 0.053 | -8.336 8.326 |
| OTU651 | - | Bacilli | 0.065 | 4.152 | 0.056 | -8.227 8.192 | OTU543 | - | Clostridia | 0.044 | 4.170 | 0.056 | -8.159 8.275 |

From the results, OTUs of belonging to the *Clostridia* and *Bacilli* class of bacteria predominate the top associations, that is, are more likely to be associated with atherosclerosis. This aligns with the literary knowledge of their central role in atherosclerosis susceptibility.

The other identified OTU classes, including *Gammproteobacteria* [22], *Actinobacteria* [23], *Clostridia* [24], *Bacilli* [25], and *Bacteroidia* [25] have also been associated with susceptibility to atherosclerosis. Thus, the associations identified by our model replicates those reported in previous literature, at least at the class level; lending support to the robustness of the estimation and the biological relevance of the associations.

## 4. Conclusion

Microbiome association studies, prodded by the advent and development of high-throughput sequencing technologies, offer unparalleled opportunities to elucidate the underlying mechanisms of complex human traits and diseases. In modelling the link between microbiome taxonomic features and the host’s phenotypic status, considerations ought to be made of the underlying dataset.

A statistical model that incorporates taxa similarity and models sample heterogeneity will fit the data better and provide more accurate measure of taxa effects. Here, we formulated a Bayesian statistical framework, phyloDPM for analysis of 16Sr RNA microbiome data. Unlike many existing methods that associates overall microbiome profile with the host’s phenotype, phyloDPM allows estimation of OTU-specific effects. Besides accounting for OTU clustering at the various phylogenetic depths, the method, importantly, models multi-collinearity among taxa as well as heterogeneity across samples. Moreover, it is robust to the possibility that the phylogenetic tree may be mis-specified or uninformative with respect to OTU effects on the host’s phenotypic state. Jointly modelling taxa similarities as well as latent heterogeneity facilitates identification of true phenotype-associated taxa and their effects. By simulation studies, we demonstrated that phyloDPM has satisfactory performance. In the real microbiome data studies of schizophrenia, and atherosclerosis, we further demonstrated the merits of the method in capturing association signals.

However, some key methodological limitations and assumptions of the framework require mention. First, the proposed model makes a key assumption that OTU effects is additive and that higher order interaction effects is negligible, which permits modelling the joint effect of OTUs as a linear combination of individual OTU effects using only main effects. Although often sufficient in practice, we expect that the inclusion of a few higher order terms could prove useful by way of improving precision on parameter estimates.

Second, the accuracy of the model diminishes as the number of taxa in the analysis gets large; albeit one way to address this could be to apply a more stringent pre-processing procedure as a way of reducing dimensionality, so that many of the low-abundant OTUs, which can be regarded as being less biologically informative and noisy, are filtered out. This must, however, be done with care; otherwise rare but phenotype-informative taxa may be erroneously discarded. Finally, the model is primarily developed for case-control phenotypes. Nevertheless, extension to quantitative phenotype can be made in a straight-forward way. The only modification would be replacing the logistic model structure with its linear counterpart. Gratifyingly, in this case, the conditional distributions of the parameters assume standard distributional forms that can be easily sampled from, and hence Gibbs sampling, with a Metropolis step for the precision parameter of the Dirichlet process component, can be directly applied.

In conclusion, the proposed approach represents a potentially robust way to model host’s OTU-phenotype relationship, especially in light of multicollinearity and the noise - from plasticity to the host’s internal and external factors - that characterise microbiome abundance data.

## Data availability

All analyses involved publicly available data that are accessible with the provided accession codes (Qiita, https://qiita.ucsd.edu; study ID 349, and study ID 11710).

## Acknowledgments

The authors thank the South African Centre for High Performance Computing (CHPC) for the computing facility that were utilized some some of the computational analyses. Responsibility for the contents of the article lies entirely with the author.

## Author contribution

D.A. and E.C.R. conceived the model for the statistical framework. D.A. developed the framework and wrote the scripts. D.A and E.C.R analysed the results. D.A drafted the manuscript. E.C.R revised and reviewed the manuscript.

