## Supplementary Table and Supplementary Method for "Modelling Microbiome Association with Host Phenotypes Using a Bayesian Dirichlet Process Model"

### 1 MATERIALS AND METHODS

#### 1.1 Dirichlet Process for the Random Effects $\psi$

The DP, by the formulation of [5], is a probability measure defined on the probability space $(\Omega, \mathcal{A}, G)$ of all measures $G$ such that for any arbitrary and finite partition $\{B_1, B_2, \cdots, B_k\}$ of $\Omega$, the joint distribution of the vector $[G(B_1), G(B_2), \cdots, G(B_k)]$ is a Dirichlet distribution with parameters $[mG_0(B_1), mG_0(B_2), \cdots, mG_0(B_k)]$, $m(> 0)$, is a concentration (precision) parameter and $G_0$ is a base distribution defined on $\Omega$. For any such measurable set $B$, we have $E[G(B)] = G_0(B)$.

The DP provides a flexible approach for modelling latent heterogeneity through the construction of probability measures on the space of distribution functions. Assume each $\psi_i$ is a realization from a general distribution that follows a DP, and suppose further that the base distribution of the DP is normal. That is,

$$\psi_i \sim G\,; \quad G \sim \mathrm{DP}(m, G_0)\,, \quad G_0 = \mathrm{N}(0, \tau^2)\,.$$

In this formalism, we postulate the random effects to be generated by a mixture of “atoms" of underlying distributions of known parametric form; Gaussian in this case.

The two elements $G_0$, the base distribution, and $m$, the concentration parameter, completely specify the DP. $G_0$ defines the probability distribution over the space of component parameters, and serves as a prior for the unknown distribution $G$. Meanwhile, $m$ defines a probability distribution on the component weights, and expresses our uncertainty about the form of $G$.

The predictive distribution of $\boldsymbol{\psi} = (\psi_1, \psi_2, \cdots, \psi_n)$ is

$$\pi(\boldsymbol{\psi}|G_0, m) = \int_G \Big[\prod_{i=1}^{n} \pi(\psi_i|G)\pi(G|G_0, m)\Big] \mathrm{d}G, \tag{1}$$

which can be decomposed as

$$\begin{aligned}\pi(\boldsymbol{\psi}|G_0, m) =& \pi(\psi_1|G_0, m) \times \pi(\psi_2|G_0, m, \psi_1) \times \cdots \times \\ & \pi(\psi_n|G_0, m, \psi_1, \cdots, \psi_{i-1}) \,.\end{aligned}$$

Following [1], we have $\psi_1|G_0, m \sim G_0$, and the successive conditional distributions are

$$\begin{aligned}\psi_i|G_0, m, \psi_1, \cdots, \psi_{i-1} \sim \frac{m}{i-1+m} G_0(\psi_i) + \\ \frac{1}{i-1+m} \sum_{j=1}^{i-1} \delta(\psi_j = \psi_i) \,,\end{aligned} \tag{2}$$

where $\delta(\cdot)$ is the usual Dirac delta function.

#### 1.2 Model Likelihood

Given $n$ independent measurements, the joint distribution of the observed data in Eqn (**??**) can be expressed in a general form as

$$\begin{aligned} y_1, \cdots, y_n &\sim f(y_1, \cdots, y_n | \boldsymbol{\theta}, \psi_1, \cdots, \psi_n) \\ &= \prod_{i=1}^{n} f(y_i | \boldsymbol{\theta}, \psi_i), \end{aligned} \tag{3}$$

where $\boldsymbol{\theta}$ is a vector of all except the DP model parameters. The likelihood function, $L(\boldsymbol{\theta}|\mathbf{y})$, can then be obtained, by definition, by integrating over the random effects. Therefore

$$\begin{aligned} L(\boldsymbol{\theta}|\mathbf{y}) = \int & f(y_1, \cdots, y_n | \boldsymbol{\theta}, \psi_1, \psi_n) \times \\ & \pi(\psi_1, \cdots, \psi_n) \mathrm{d}\psi_1 \cdots \mathrm{d}\psi_n \,, \end{aligned} \tag{4}$$

where $\pi(\psi_1, \cdots, \psi_n) = \pi(\boldsymbol{\psi})$ is as previously defined in Eqn (1).

The clustering in the DP model leads to a succinct mathematical formalism in which the observations can be represented by a binary matrix. Following [7, 8], let $A$ be a $n \times k$ binary matrix associated with a particular partition of the sample of size $n$ into $k$ groups, $k = 1, \cdots, n$. A partition as such is a cluster or, in a more appropriate terminology, a "subcluster" $C$, since the grouping is done non-parametrically and the resulting clusters may differ from that determined by some substantive criteria. The matrix $A$ has the general form,

$$A = \begin{pmatrix} a_{11} & a_{12} & \cdots & a_{1k} \\ a_{21} & a_{22} & \cdots & a_{2k} \\ \vdots & & \cdots & \vdots \\ a_{n1} & a_{n2} & \cdots & a_{nk} \end{pmatrix}$$

Each row of $A$ has all zeros except for a 1 in one position indicating the group the observation belongs. Meanwhile, the column sums of $A$ represent the number of observations in each of $k$ groups. This means that, using the matrix $A$, the $n$-dimensional vector $\boldsymbol{\psi}$ can be represented by a $k(\leq n)$-dimensional vector $\boldsymbol{\eta}$. Then $\psi_i = \eta_j$ so that $\boldsymbol{\psi} = A\boldsymbol{\eta}$, where $\boldsymbol{\eta} \sim N_k(0, \tau^2 I_k)$.

The likelihood function, the joint density of the observations as a function of the parameters, becomes

$$
\begin{aligned}
& L_k(\boldsymbol{\theta}|\mathbf{y}) = f(\mathbf{y}|\boldsymbol{\theta}) \\
&= \int \prod_{i=1}^{n} \Big[\frac{1}{1+\exp\{-\big(X_i'\boldsymbol{\alpha} + Z_i'\boldsymbol{\beta} + (A\boldsymbol{\eta})_i\big)\}}\Big]^{y_i} \\
& \Big[\frac{1}{1+\exp\{X_i'\boldsymbol{\alpha} + Z_i'\boldsymbol{\beta} + (A\boldsymbol{\eta})_i\}}\Big]^{1-y_i} \mathrm{d}G_0(\boldsymbol{\eta}) \\
&= \prod_{i=1}^{n} \Big[\frac{1}{1+\exp\{-\big(X_i'\boldsymbol{\alpha} + Z_i'\boldsymbol{\beta} + (A\boldsymbol{\eta})_i\big)\}}\Big]^{y_i} \\
& \Big[\frac{1}{1+\exp\big\{X_i'\boldsymbol{\alpha} + Z_i'\boldsymbol{\beta} + (A\boldsymbol{\eta})_i\big\}}\Big]^{1-y_i} \times \\
& \prod_{j=1}^{k} \left(\frac{1}{2\pi\tau^2}\right)^{\frac{1}{2}} \exp\left(-\frac{1}{2\tau^2}\eta_j^2\right). \qquad (5)
\end{aligned}
$$

To complete the Bayesian modelling, we specify the following priors on the other model parameters:

$$
\begin{aligned}
& \boldsymbol{\alpha}|\sigma_\alpha^2 \sim N_q(\mathbf{0}, \sigma_\alpha^2 I_q), \quad \tau^2 \sim \mathrm{IG}(a_\tau, b_\tau) \\
& \sigma_\alpha^2 \sim \mathrm{IG}(a_\alpha, b_\alpha), \text{ and } \sigma_\beta^2 \sim \mathrm{IG}(a_\beta, b_\beta) \ . \qquad (6)
\end{aligned}
$$

#### 1.3 Bayesian Inference of the Parameters

In this section, we consider estimation of the parameters involved in our Dirichlet process logistic random effects model. The ensemble of parameters can be considered into three sets: (i) $A$, the matrix of sub-clusters (ii) $m$, the precision parameter of the Dirichlet process, and (iii) $\boldsymbol{\theta} = (\boldsymbol{\alpha}, \boldsymbol{\beta}, \boldsymbol{\mu}, \boldsymbol{\gamma}, \tau_\gamma^2, \boldsymbol{\eta}, \tau^2, \sigma_\alpha^2, \sigma_\beta^2)$, the other model parameters. Using the algorithmic idea in [6], these three sets of parameters can be iteratively estimated group-wise as follows: conditional on $A$

and $m$, simulate $\boldsymbol{\theta}$; then conditional on $\boldsymbol{\theta}$ and $A$, simulate $m$; finally, conditional on $\boldsymbol{\theta}$ and $m$, simulate $A$.

#### 1.4 Estimation of Model Parameters, $\theta$

Given a parameter $\boldsymbol{\theta}$ and a set of observed data $y$, the Bayesian approach seeks to obtain the probability of the parameter $\boldsymbol{\theta}$ given the set of data available $y$, mathematically expressed as $p(\boldsymbol{\theta}|y)$. Given a prior information about the parameter $\boldsymbol{\theta}$, which is expressed as a probability distribution $p(\boldsymbol{\theta})$. The interest is in the posterior distribution of the parameter given then data, $p(\boldsymbol{\theta}|y)$. This posterior distribution is obtained by combining the prior distribution with the likelihood function, using Baye's theorem. Applying this theorem, the likelihood in Eqn 5 can be combined with the priors in Eq 6 to give the following joint posterior distribution.

$$\begin{aligned} f(\boldsymbol{\theta}|A,\mathbf{y}) \propto L(\boldsymbol{\theta}|A,\mathbf{y}) \times \left(\frac{1}{\tau^2}\right)^{a_\tau+1} \exp\left(-\frac{b_\tau}{\tau^2}\right) \times \\ \left(\frac{1}{\sigma_\alpha^2}\right)^{\frac{q}{2}} \exp\left(-\frac{|\boldsymbol{\alpha}|^2}{2\sigma_\alpha^2}\right) \times \left(\frac{1}{\sigma_\beta^2}\right)^{\frac{p}{2}} \exp\left(-\frac{(\boldsymbol{\beta}-\boldsymbol{\mu})^2}{2\sigma_\beta^2}\right) \times \\ \left(\frac{1}{\sigma_\alpha^2}\right)^{a_\alpha+1} \exp\left(-\frac{b_\alpha}{\sigma_\alpha^2}\right) \times \left(\frac{1}{\sigma_\beta^2}\right)^{a_\beta+1} \exp\left(-\frac{b_\beta}{\sigma_\beta^2}\right) \times \\ \exp\left(-\frac{(\boldsymbol{\mu}-\boldsymbol{\gamma})C_T^{-1}(\boldsymbol{\mu}-\boldsymbol{\gamma})}{2}\right) \times \exp\left(-\frac{|\boldsymbol{\gamma}|^2}{2\tau_\gamma^2}\right) . \end{aligned} \tag{7}$$

#### 1.5 Full Conditional Distributions

We now proceed to derive the full-conditional distribution for each parameter in the model. For notational convenience, let $\boldsymbol{\theta}$ be the vector of all model parameters and $\boldsymbol{\theta}_{-p}$ be the vector of all these parameters except $p$. Let

$$p_i = P\left(y_i = 1|\boldsymbol{\theta}, X, Z, \mathbf{y}\right) = \frac{\exp\left\{X_i'\boldsymbol{\alpha} + Z_i'\boldsymbol{\beta} + (A\boldsymbol{\eta})_i\right\}}{1 + \exp\left\{X_i'\boldsymbol{\alpha} + Z_i'\boldsymbol{\beta} + (A\boldsymbol{\eta})_i\right\}} \tag{8}$$

and let $q_i = 1 - p_i$.

For each response variable $y_i$, we introduce two mutually independent latent variables $\lambda_i$ and $\phi_i$, where each is assumed to be uniformly distributed over the interval $(0,\ 1)$. Then, according to Eqn 8, the variables $\lambda_i$ and $\phi_i$ are constrained by the response variable $y_i$. When $y_i = 1$, we have $p_i = \mathrm{E}_\lambda\left[I\left(0 < \lambda_i \leq p_i\right)\right]$, and when $y_i = 0$, we have $q_i = \mathrm{E}_{\phi_i}\left[I\left(0 < \phi_i \leq q_i\right)\right]$. Therefore, the likelihood function in Eqn 5 can be rewritten using the expectation of the auxiliary variables. That is,

$$L(\boldsymbol{\theta}|A, \mathbf{y}) = \prod_{i=1}^{n}\left[I(y_i = 1)I(0 < \lambda_i \leq p_i) + I(y_i = 0)I(0 < \phi_i \leq q_i)\right] \times \prod_{j=1}^{k}\left(\frac{1}{2\pi\tau^2}\right)^{\frac{1}{2}}\exp\left(-\frac{1}{2\tau^2}\eta_j^2\right). \tag{9}$$

Using (7) and (9), we obtain the following full conditional distributions:

**(a)** Full conditionals for $\lambda$ and $\phi$ are both uniform.
For $i = 1, \cdots, n$, we have

$$\lambda_i|\boldsymbol{\theta}_{-\lambda_i}, X, Z, \mathbf{y} \sim \text{Uniform}\left(0,\ p_i\right), \quad \text{if } y_i = 1 \tag{10}$$

$$\phi_i|\boldsymbol{\theta}_{-\phi_i}, X, Z, \mathbf{y} \sim \text{Uniform}\left(0,\ q_i\right), \quad \text{if } y_i = 0 \tag{11}$$

The full conditional distribution for $\boldsymbol{\alpha}$, $\boldsymbol{\beta}$ and $\boldsymbol{\eta}$ are, respectively, the following truncated distributions:

**(b)** Full conditional distribution for $\alpha_j;\ j \in \{1, \cdots, q\}$:

Let $S_1 = \{(i,j) : y_i = 1, X_{ij} > 0\}$ and $S_2 = \{(i,j) : y_i = 0, X_{ij} > 0\}$, then we have

$$\alpha_j | \boldsymbol{\theta}_{-\alpha_j}, X, Z, \mathbf{y} \sim \begin{cases} N(0, \sigma_\alpha^2)\,, & \alpha_j \in \left[\alpha_j^L, \alpha_j^U\right] \\ 0\,, & \text{otherwise} \end{cases} \tag{12}$$

where

$$\alpha_j^L = \max_{i \in S_1} \left\{ \frac{1}{X_{ij}} \left[ \log\left( \frac{\lambda_i}{1-\lambda_i} \right) - \sum_{k \neq j} X_{ik}\alpha_k - Z_i'\boldsymbol{\beta} - (A\boldsymbol{\eta})_i \right] \right\} \quad \text{and}$$

$$\alpha_j^U = \min_{i \in S_2} \left\{ \frac{1}{X_{ij}} \left[ \log\left( \frac{1-\phi_i}{\phi_i} \right) - \sum_{k \neq j} X_{ik}\alpha_k - Z_i'\boldsymbol{\beta} - (A\boldsymbol{\eta})_i \right] \right\}.$$

**(c)** Full conditional distribution for $\beta_j; j \in \{1, \cdots, p\}$. Because some elements of the matrix $L$ can be negative, let $S_1 = \{(i,j) : y_i = 1, Z_{ij} > 0\}$, $S_2 = \{(i,j) : y_i = 0, Z_{ij} < 0\}$, $S_3 = \{(i,j) : y_i = 1, Z_{ij} < 0\}$, and $S_4 = \{(i,j) : y_i = 0, Z_{ij} > 0\}$. Then, we have

$$\beta_j | \boldsymbol{\theta}_{-\beta_j}, X, Z, \mathbf{y} \sim \begin{cases} N(\mu_j, \sigma_\beta^2)\,, & \beta_j \in \left[\beta_j^L, \beta_j^U\right] \\ 0\,, & \text{otherwise} \end{cases} \tag{13}$$

where

$$\beta_j^L = \max\left\{\max_{i\in S_1}\left\{\frac{1}{Z_{ij}}\left[\log\left(\frac{\lambda_i}{1-\lambda_i}\right) - X_i'\boldsymbol{\alpha} - \sum_{k\neq j} Z_{ik}\beta_k - (A\boldsymbol{\eta})_i\right]\right\}, \max_{i\in S_2}\left\{\frac{1}{Z_{ij}}\left[\log\left(\frac{1-\phi_i}{\phi_i}\right) - X_i'\boldsymbol{\alpha} - \sum_{k\neq j} Z_{ik}\beta_k - (A\boldsymbol{\eta})_i\right]\right\}\right\}$$

and

$$\beta_j^U = \min\left\{\min_{i\in S_3}\left\{\frac{1}{Z_{ij}}\left[\log\left(\frac{\lambda_i}{1-\lambda_i}\right) - X_i'\boldsymbol{\alpha} - \sum_{k\neq j} Z_{ik}\beta_k - (A\boldsymbol{\eta})_i\right]\right\}, \min_{i\in S_4}\left\{\frac{1}{Z_{ij}}\left[\log\left(\frac{1-\phi_i}{\phi_i}\right) - X_i'\boldsymbol{\alpha} - \sum_{k\neq j} Z_{ik}\beta_k - (A\boldsymbol{\eta})_i\right]\right\}\right\}.$$

**(d)** Full conditional distribution for $\eta_k$; $k \in \{1, \cdots, K\}$. Let $S_1 = \{i : y_i = 1\}$ and $S_2 = \{i : y_i = 0\}$, then we have

$$\eta_k|\boldsymbol{\theta}_{-\eta_k}, X, Z, \mathbf{y} \sim \begin{cases} N(0,\tau^2)\,, & \eta_k \in \left[\eta_k^L, \eta_k^U\right] \\ 0\,, & \text{otherwise} \end{cases} \tag{14}$$

where $\eta_k^L = \max_{S_1}\left\{\log\left(\frac{\lambda_i}{1-\lambda_i}\right) - X_i'\boldsymbol{\alpha} - Z_i'\boldsymbol{\beta}\right\}$, and $\eta_k^U = \min_{S_2}\left\{\log\left(\frac{1-\phi_i}{\phi_i}\right) - X_i'\boldsymbol{\alpha} - Z_i'\boldsymbol{\beta}\right\}$.

**(e)** The full conditional distribution for $\boldsymbol{\mu}$.

$$f(\boldsymbol{\mu}|\cdots) \propto$$
$$\exp\left\{-\frac{1}{2}\left[\frac{1}{\sigma_\beta^2}(\boldsymbol{\beta}-\boldsymbol{\mu})^2+(\boldsymbol{\mu}-\boldsymbol{\gamma})C_T^{-1}(\boldsymbol{\mu}-\boldsymbol{\gamma})\right]\right\} \tag{15}$$

The quadratic forms of $\boldsymbol{\mu}$ in Eqn (15) can be combined using the **(author?)** [3] identity for quadratic functions. The identity states that

$$\begin{aligned}
&(\mathbf{z}-\mathbf{m})'\mathbf{M}(\mathbf{z}-\mathbf{m})+(\mathbf{z}-\mathbf{b})'\mathbf{B}(\mathbf{z}-\mathbf{b}) = \\
&(\mathbf{z}-\mathbf{c})'(\mathbf{M}+\mathbf{B})(\mathbf{z}-\mathbf{c})+ \\
&(\mathbf{m}-\mathbf{b})'\mathbf{M}(\mathbf{M}+\mathbf{B})'\mathbf{B}(\mathbf{m}-\mathbf{b})\,,
\end{aligned} \tag{16}$$

where $\mathbf{c}=(\mathbf{M}+\mathbf{B})^{-1}(\mathbf{Mm}+\mathbf{Bb})$.

Using this identity, we can combine the sums in Eqn (15) as

$$\begin{aligned}
&\frac{1}{\sigma_\beta^2}(\boldsymbol{\beta}-\boldsymbol{\mu})^2+(\boldsymbol{\mu}-\boldsymbol{\gamma})C_T^{-1}(\boldsymbol{\mu}-\boldsymbol{\gamma}) = \\
&(\boldsymbol{\mu}-\hat{\boldsymbol{\mu}})'\left(\frac{1}{\sigma_\beta^2}I+C^{-1}\right)(\boldsymbol{\mu}-\hat{\boldsymbol{\mu}})+ \\
&(\boldsymbol{\beta}-\boldsymbol{\gamma})'\frac{1}{\sigma_\beta^2}\left(\frac{1}{\sigma_\beta^2}I+C^{-1}\right)^{-1}C^{-1}(\boldsymbol{\beta}-\boldsymbol{\gamma})\,,
\end{aligned} \tag{17}$$

where $\hat{\boldsymbol{\mu}}=\left(\frac{1}{\sigma_\beta^2}I+C^{-1}\right)^{-1}\left(\frac{1}{\sigma_\beta^2}\boldsymbol{\beta}+C^{-1}\boldsymbol{\gamma}\right)$.

Putting Eqn (17) in Eqn (15) and retaining only the parts that varies with $\boldsymbol{\mu}$, the full conditional distribution can be expressed as

$$f(\boldsymbol{\mu}|\cdots) \propto \exp\left\{-\frac{1}{2}(\boldsymbol{\mu}-\hat{\boldsymbol{\mu}})\left(\frac{1}{\sigma_\beta^2}I+C^{-1}\right)(\boldsymbol{\mu}-\hat{\boldsymbol{\mu}})\right\}\,,$$

which is immediately recognized as the kernel of a Gaussian distribution with mean vector $\hat{\boldsymbol{\mu}}$, as defined above, and variance-covariance matrix $\left(\frac{1}{\sigma_\beta^2} I + C^{-1}\right)$. Thus, component-wise, we can write

$$\mu_j | \cdots \sim \mathcal{N}\left(\left\{\left(\frac{1}{\sigma_\beta^2} + (C_j' C_j)^{-1}\right)^{-1}, \right.\right.$$
$$\left.\left.\left(\frac{1}{\sigma_\beta^2}\beta_j + (C_j' C_j)^{-1}\gamma_j\right)\right\}\left(\frac{1}{\sigma_\beta^2} + (C_j' C_j)^{-1}\right)\right)$$

**(f)** The full conditional distribution for $\boldsymbol{\gamma}$.

Following similar exposition of part **(e)** above, we find that

$$\gamma_j | \cdots \sim \mathcal{N}\left(\left\{\left(\frac{1}{\tau_\gamma^2} + (C_j' C_j)^{-1}\right)^{-1}\left((C_j' C_j)^{-1}\mu_j\right)\right\}, \right.$$
$$\left.\left(\frac{1}{\tau_\gamma^2} + (C_j' C_j)^{-1}\right)\right)$$

**(g)** Finally, the full conditional distribution for $\tau^2$, $\sigma_\alpha^2$, and $\sigma_\beta^2$ are, respectively:

$$\tau^2 | \boldsymbol{\theta}_{-\tau^2}, X, Z, \mathbf{y} \sim \text{IG}\left(\frac{k}{2} + a_\tau, \frac{1}{2}|\boldsymbol{\eta}|^2 + b_\tau\right) \tag{18}$$

$$\sigma_\alpha^2 | \boldsymbol{\theta}_{-\sigma_\alpha^2}, X, Z, \mathbf{y} \sim \text{IG}\left(\frac{q}{2} + a_\alpha, \frac{1}{2}|\boldsymbol{\alpha}|^2 + b_\alpha\right) \tag{19}$$

$$\sigma_\beta^2 | \boldsymbol{\theta}_{-\sigma_\beta^2}, X, Z, \mathbf{y} \sim \text{IG}\left(\frac{p}{2} + a_\beta, \frac{1}{2}(\boldsymbol{\beta} - \boldsymbol{\mu})^2 + b_\beta\right), \tag{20}$$

where $\text{IG}(\cdot, \cdot)$ denotes the inverse Gamma distribution.

#### 1.6 Generation of Sub-clusters Matrix $A$

For the generation of the sub-clusters matrix $A$, we apply the method of [6], where a Metropolis-Hastings algorithm with a Dirichlet proposal distribution is used. The method is based on recognizing the fact that the marginal distribution of any $k$ components taken from an $n$-dimensional multinomial/Dirichlet distribution is also multinomial/Dirichlet. Therefore, a sampling scheme can be employed with candidate draws taken from a multinomial/Dirichlet distribution.

Starting from the model parameters at iteration $t$:
$\boldsymbol{\theta}^{(t)} = \left(\boldsymbol{\alpha}^{(t)}, \boldsymbol{\beta}^{(t)}, \boldsymbol{\mu}^{(t)}, \boldsymbol{\gamma}^{(t)}, \tau_{\gamma}^{2(t)}, \boldsymbol{\eta}^{(t)}, \tau^{2(t)}, \sigma_{\alpha}^{2(t)}, \sigma_{\beta}^{2(t)}\right)$ and $A^{(t)}$, the Dirichlet distribution parameters is first simulated from:

$$\begin{aligned}\boldsymbol{q}^{(t+1)} &= (q_1^{(t+1)}, q_2^{(t+1)}, q_n^{(t+1)}) \\ &\sim \text{Dirichlet}(s_1^{(t)} + 1, s_2^{(t)} + 1, \cdots, s_k^{(t)} + 1, \overbrace{1, 1, 1, \cdots, 1}^{n-k \text{ components}}),\end{aligned}$$

where $s_j \geq 1$ are the non-zero column sums of $A$, and $s_1 + s_2 + \cdots + s_k = n$. Then based on the value of $\boldsymbol{q}^{(t+1)}$, we generate $A^{(t+1)}$ from a Multinomial distribution,

$$\begin{aligned}A^{(t+1)} \sim & P(A^{(t)}) f(y|\theta^{(t+1)}, A^{(t)}) \\ & \binom{n}{n_1, \cdots, n_{k'}} \prod_{j=1}^{k'} \left[q_j^{(t+1)}\right]^{n'_j}.\end{aligned}$$

The columns with zero column sums are then deleted, resulting in a $n \times k$ matrix. Deleting the zero sum columns is a marginalization of the multinomial distribution.

### 2 APPLICATION

#### 2.1 Simulating OTU Abundance

OTU counts were simulated using a DM distribution. In order to make the simulation more realistic, the parameters of the DM distribution (mean and dispersion) were estimated from real microbiome data of human upper respiratory microbiome in the study by [4]. This dataset contains the counts of 856 OTUs from 60 samples. Also available from this dataset is the phylogenetic tree structure for these OTUs.

The $p \in \{200, 400, 800\}$ most abundant OTUs from this dataset were used for parameter estimation, and, accordingly, $p$ OTUs were subsequently simulated for $n = 500$ individuals. The simulated OTU counts data closely mimic real data (**Figure ??**), characterized by zero-inflation and overdispersion.

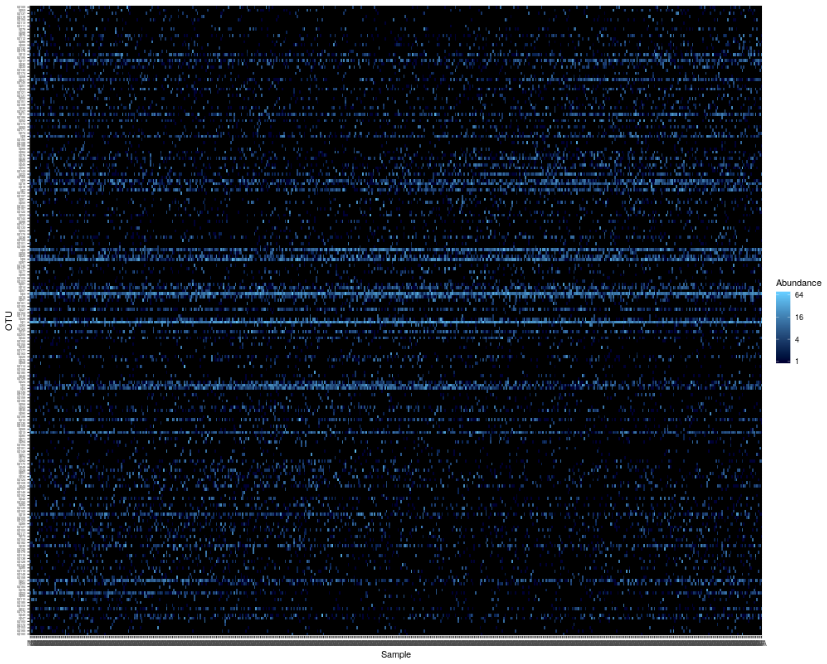


(a)

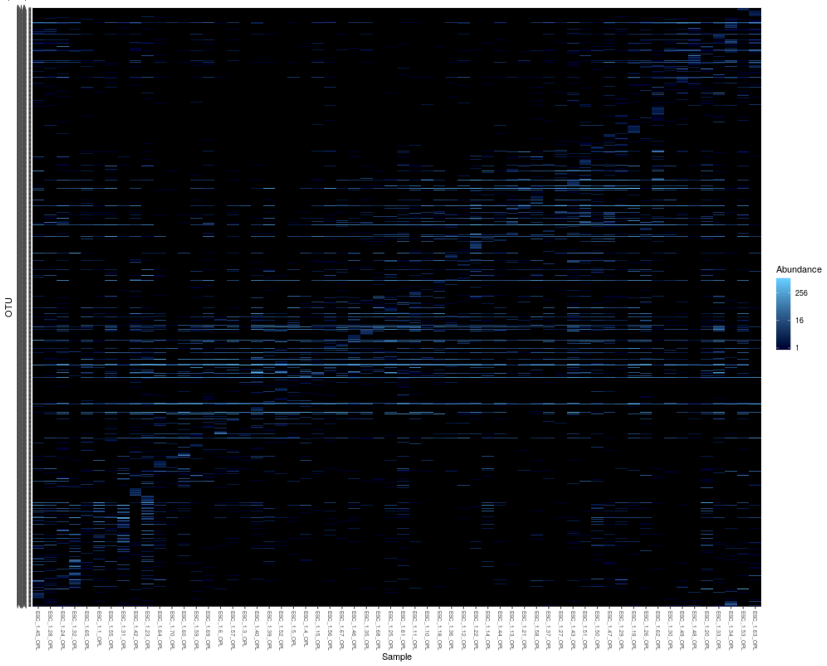


(b)

Supplementary Figure 1: Heatmap showing the OTU abundance distribution for (a) simulated microbial community from 500 samples, and (b) real microbial community of the human upper respiratory tract from 60 samples. The scale indicates abundance level of the OTU counts with dark representing zero counts (see legends).

The resulting OTU count data for each sample was normalized by the total read count for that sample, where the total read count for a sample was obtained by sampling from a negative binomial distribution with mean 5000 and dispersion 25, which reflects typical sequencing depths in empirical microbiome studies.

#### 2.2 Choice of Hyperparameters and Sensitivity Analysis

In applying the model to the datasets, the hyper-parameters for the Bayesian priors were set as follows. Following the simulation study by [10], we let the parameters $c$ and $d$ for the Gamma distribution of the precision parameter $m \sim \Gamma(c, d)$ be $c = d = \exp(-0.033 \times n)$ where $n$ is the sample size. For the variance components, while the use of weakly informative inverse gamma hyper-parameters such as $\Gamma(0.1, 0.1)$ or $\Gamma(1, 0.001)$ is popular, our test simulations showed their use leads to very poor MCMC posterior samples; possibly given that such hyperparameters place the majority of the prior mass away from zero. Therefore, it is imperative to carry out a sensitivity analysis to assess the impact of hyper-parameter values on the performance of the model, and, subsequently, to specify appropriate hyper-parameter values. To conduct sensitivity analysis with respect to the hyper-parameter values, we performed several simulation runs, varying hyper-parameter values $(a_\tau,\, b\mathcal{T})$ and $(a_\beta,\, b_\beta)$ from 1 to 20, and used Matthews correlation coefficient (MCC) [9] to compare the results from the different simulations.

Specifically, we first conduct initial test runs with hyper-parameter values $(a_\tau,\, b\mathcal{T})$ and $(a_\beta,\, b_\beta)$ from 1 to 20, monitoring the Bayesian posterior distribution of the parameters. Based on this, $(a_\tau,\, b\mathcal{T})$ is fixed to a value $(a^*_\tau,\, b^*_\mathcal{T}) \in \{1,\, 20\}$ that provide stable posterior estimates, and several simulation runs are performed for various values of $(a_\beta,\, b_\beta)$ from 1 to 20. Then, in each replicate, the discriminatory power of the model (in discerning case from control subjects) is evaluated by calculating the number of true positive (TP), true negative (TN), false positive

(FP) and false negative (FN), from where MCC is calculated as

$$\text{MCC} = \frac{\text{TP} \times \text{TN} - \text{FP} \times \text{FN}}{\sqrt{(\text{TP}+\text{FP})(\text{TP}+\text{FN})(\text{TN}+\text{FP})(\text{TN}+\text{FN})}}\,.$$

MCC score takes values in the interval $[-1, 1]$, with $1$ showing a complete agreement (each case subject is classified as a case and each control subject is classified as a control), and $-1$ a complete disagreement (a case subject is classified as control and *vice versa*). MCC provides a good measure of the global accuracy, since it accounts for both sensitivity and specificity. Because of this, it is widely used as a performance metric in Bioinformatics [2]. Following the above procedure, we take the 'appropriate' hyper-parameter pair $(a_\beta,\, b_\beta)$ to be the pair that yields the highest value of the MCC by choosing, out of all the tested hyper-parameter settings. Similarly, we determine the 'appropriate' hyper-parameter $(a_\tau,\, b\tau)$ by fixing $(a_\beta,\, b_\beta)$ at the obtained optimal value, and varying $(a_\tau,\, b\tau)$ over the same interval (1 to 20) and calculating MCC for several replicates. Figure ?? displays the heat-map of average MCC values with respect to various choices of inverse Gamma hyper-parameters. As can be observed, for $\tau^2$, the MCC is undesirable for values of $a_\tau \leq 1$ and $b_\tau \leq 1$. Similarly, for $\sigma^2_\beta$, MCC is sub-optimal for $a_\beta \leq 5$ and $b_\beta \leq 5$.

Meanwhile, fixing one of hyper-parameter values $a_\tau$ ($a_\beta$ for $\sigma^2_\beta$) or $b_\tau$ ($b_\beta$ for $\sigma^2_\beta$) to a small value and increasing the other component still result in a poor posterior, possibly since the associated IG($a$, $b$) prior with mean $b/(a-2)$ and variance $b^2/(a-2)^2$ is strongly informative in these scenarios. We complemented the sensitivity analysis by monitoring the changes in Bayesian posterior estimates of the parameters, on a grid of values of the hyper-parameters in the same interval (1 to 20). From these analyses, we chose $a_\tau = b_\tau = 20$ and $a_\beta = 20\,, b_\beta = 15$ as the well performing hyper-parameter configurations for the variance parameters $\sigma^2_\beta$ and $\tau^2$ respectively.

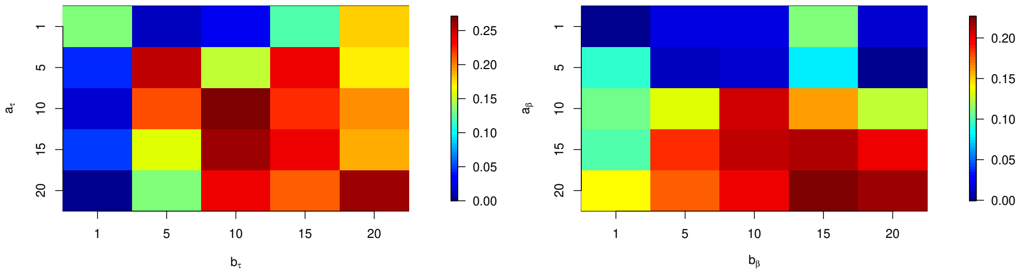


Supplementary Figure 2: Heatmap showing Matthews correlation coefficient (MCC) values for various choices of $(a, b)$ hyper-parameters for the inverse-gamma prior on the variance components tau2 and sigma2. Each MCC value is the average obtained for 10 replications.

Meanwhile, the kernel coefficient of the RBF is fixed to $10^{-3}$ for all analyses. This parameter setting achieved optimal performance in test runs. Each simulation experiment was conducted for 50 replications.

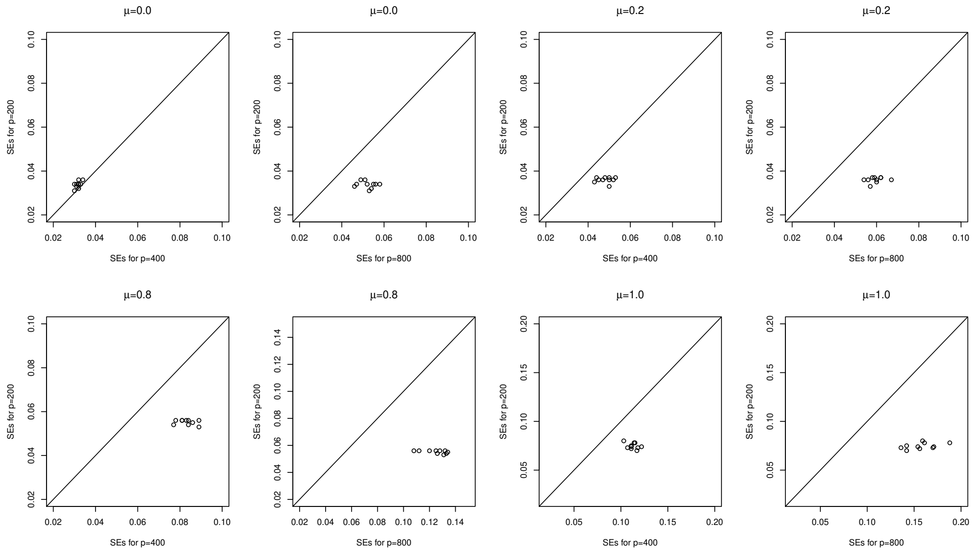

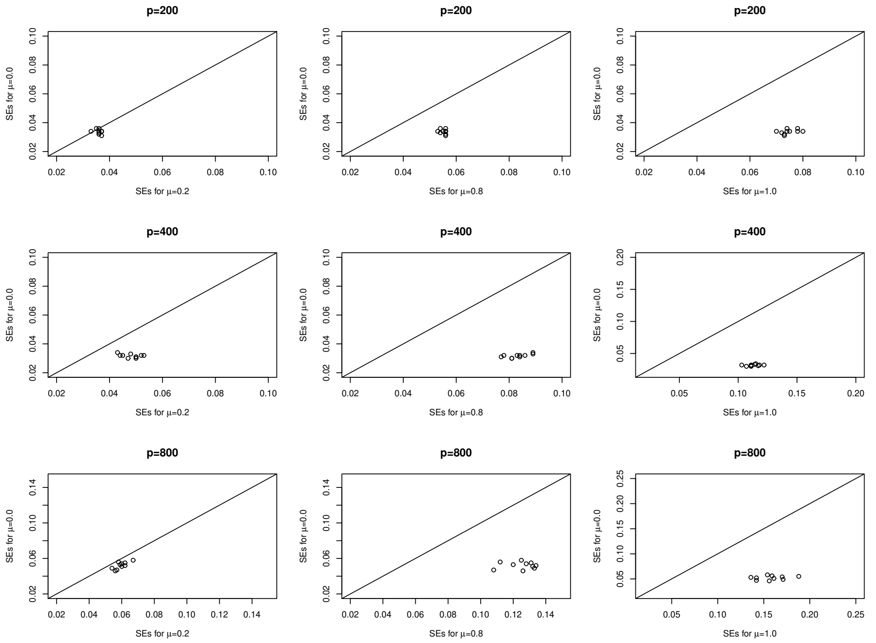


Supplementary Figure 3: Comparison of standard errors (SEs) of the model for fixed $p$ and varying $\mu$.

[6] Minjung Kyung, Jeff Gill, and George Casella. Sampling schemes for generalized linear dirichlet random effects models. Technical report, Technical Report, University of Florida, Department of Statistics. Available ..., 2009.

[7] Minjung Kyung, Jeff Gill, George Casella, et al. Estimation in dirichlet random effects models. *The Annals of Statistics*, 38(2):979–1009, 2010.

[8] Chen Li, George Casella, and Malay Ghosh. Estimation of regression vectors in linear mixed models with dirichlet process random effects. *Communications in Statistics-Theory and Methods*, 47(16):3935–3954, 2018.

[9] Brian W Matthews. Comparison of the predicted and observed secondary structure of t4 phage lysozyme. *Biochimica et Biophysica Acta (BBA)-Protein Structure*, 405(2):442–451, 1975.

[10] Siva Murugiah and Trevor Sweeting. Selecting the precision parameter prior in dirichlet process mixture models. *Journal of Statistical Planning and Inference*, 142(7):1947–1959, 2012.
